# Time-kill kinetics reveal heterogeneous tolerance to disinfectants

**DOI:** 10.1101/2022.06.22.497202

**Authors:** Niclas Nordholt, Dominique Lewerenz, Frank Schreiber

**Affiliations:** Division of Biodeterioration and Reference Organisms (4.1), Department of Materials and the Environment, Federal Institute for Materials Research and Testing (BAM), Berlin, Germany

## Abstract

Disinfection is an important strategy to limit the spread of infections. Failure of disinfection may facilitate evolution of resistance against disinfectants and antibiotics through the processes of cross-resistance and co-resistance. The best possible outcome of disinfection minimizes the number of surviving bacteria and the chance for resistance evolution. Resistance describes the ability to grow in previously inhibitory concentrations of an antimicrobial, whereas tolerance is associated with enhanced survival of lethal doses. Individual bacteria from the same population can display considerable heterogeneity in their ability to survive treatment (i.e. tolerance) with antimicrobials, which can result in unexpected treatment failure. Here, we investigated how phenotypic heterogeneity affects the ability of *E. coli* to survive treatment with six different substances commonly used as active substances in disinfectants, preservatives and antiseptics. A mathematical model which assumes that phenotypic heterogeneity underlies the observed disinfection kinetics was used to infer whether time-kill kinetics were caused by a tolerant subpopulation. The analysis identified bimodal kill kinetics for benzalkonium chloride (BAC), didecyldimethylammonium chloride (DDAC), and isopropanol (Iso). In contrast, kill kinetics by chlorhexidine (CHX), glutaraldehyde (GTA), and hydrogen peroxide (H_2_O_2_) were best explained by unimodal kill kinetics underpinned by a broad distribution of tolerance times for CHX as opposed to a narrow distribution of tolerance times for GTA and H_2_O_2_. These findings have implications for the risk of disinfection failure, with potential consequences for the evolution of antimicrobial resistance and tolerance.

## Introduction

The application of disinfectants is an important strategy to control the growth of microbes, prevent the emergence of microbial infections and their spread between hosts. They are an indispensable tool to battle the ongoing antimicrobial resistance crisis (Murray et al. 2022). However, incomplete or inefficient disinfection poses the risk to facilitate the selection and evolution of bacteria that are less susceptible to disinfectants and antibiotics (Nordholt et al. 2021; Bore et al. 2007). Unnecessary or incorrect use of disinfectants thus threatens the efficacy of antimicrobials as a whole (Kampf 2018; SCENIHR 2009), especially considering their massive use on a global scale (European Commission Environment Directorate-General 2009). Using disinfectants too sparingly provokes the survival and selection of resistance at the application site. In contrast, overabundant use increases the chance of dissemination into the environment and off-site resistance evolution. To optimize the application of disinfectants it is crucial to understand their interaction with bacteria and the factors that interfere with their efficacy (García and Cabo 2018).

One important factor that determines the efficacy of antimicrobial compounds (e.g. disinfectants and antibiotics) is phenotypic heterogeneity displayed by bacterial populations (van Boxtel et al. 2017; Ackermann 2015). The heterogeneity within a bacterial population for a given trait is characterized by the distribution of trait values of individuals within the population. Traits such as cell size at division, growth rate in exponentially growing cells or metabolic activity, often give rise to a unimodal distribution (Van Heerden et al. 2017; Schreiber et al. 2016; Schreiber and Ackermann 2020). In these cases, the differences between individual cells are of a quantitative nature and emerge from a single population. However, there are also traits exhibiting bi- or multimodal distributions, suggesting qualitative differences between the emerging phenotypic subpopulations (Choi et al. 2008; Dubnau and Losick 2006; Ronin et al. 2017). Persister cells constitute one such distinct phenotype characterized by one or more subpopulations exhibiting increased tolerance (Balaban et al. 2004). In this context, tolerance is characterized by the duration for which a bacterial population can survive lethal doses of antimicrobials (Brauner et al. 2016). High levels of tolerance, as exhibited by persisters, can impair outcomes of antibiotic treatments and can facilitate the evolution of resistance and tolerance to antibiotics (Van den Bergh et al. 2016; Levin-Reisman et al. 2017; Windels et al. 2019). Recently, we showed that *E. coli* populations also harbor persister subpopulations to the widely used disinfectant benzalkonium chloride, and that repeated application failure of benzalkonium chloride resulted in the evolution of tolerance linked to changes in antibiotic susceptibility (Nordholt et al. 2021). However, we are currently lacking information about persistence to commonly used disinfectants from a variety of substance classes. This information will be important to devise application guidelines that consider the effect of heterogeneity on the efficacy of disinfectants, with the ultimate goal to minimize the evolution of tolerance and resistance to disinfectants and antibiotics.

Here, we used time kill experiments to survey the potential of *E. coli* to survive lethal stress of active biocidal substances used mainly as disinfectants, but which also find application as biocides for preservation or in medical products such as antiseptics. The aim was to identify whether heterogeneous tolerance, including the formation of tolerant subpopulations, was underlying survival. *Escherichia coli* was selected for this study because it is a model bacterium for antimicrobial persistence (Balaban et al. 2004; Levin-Reisman et al. 2017; Cameron et al. 2018; Dörr, Lewis, and Vulić 2009) and its antimicrobial resistance evolution is highly relevant (Murray et al. 2022). We used a concept from physical inactivation of microbes to quantitatively analyze the results. The underlying model allows to represent time-kill kinetics as direct measurements of the heterogeneity of tolerance times within bacterial populations (Withell 1942; Zafeiro Aspridou et al. 2019; Coroller et al. 2006). More specifically, time kill experiments can be understood as the empirical survival function which gives the probability that a cell survives for a certain time. This perspective highlights the inter-individual variability in the ability of bacteria to withstand disinfection and allows the quantification of the heterogeneity of individual cells based on a population-level readout. Here, we use a phenomenological mathematical model based on two mixed Weibull distributions to interpret disinfection kinetics (Coroller et al. 2006; Peleg and Cole 1998). The model can be parameterized to capture a wide range of shapes often observed in inactivation studies, allowing to distinguish unimodal kill kinetics of phenotype distributions of different magnitude from distributions emerging from distinct persister phenotypes. We combine these conceptual and analytical approaches with experimental evidence to characterize the extent to which phenotypic heterogeneity affects disinfection kinetics and to identify substances with a potentially increased risk of disinfection failure and resistance evolution due to the presence of tolerant subpopulations.

## Materials and Methods

### Strains and Growth conditions

The experiments were performed with *E. coli* K12 MG1655 (Blattner et al. 1997), cultured in LB Lennox medium (L3022, Sigma Aldrich; final concentrations: 10 g/L tryptone, 5 g/L yeast extract, 5 g/L NaCl). Pre-cultures were inoculated from a single-use −80°C freezer stock to a defined density of 10 cfu/ml (cfu - colony forming units) and incubated at 37°C with agitation at 220 rpm for 24 hours to stationary phase. Close attention was paid to keep the initial cell density and incubation times the same between experiments to ensure that cells reached a comparable physiology (Harms et al. 2017). The low initial cell density was chosen to avoid carryover of persisters from the stock and to allow as many generations of growth as possible (Balaban et al. 2004; Harms et al. 2017).

### Determination of MIC and MBC

Minimum inhibitory concentration (MIC) and minimum bactericidal concentration (MBC) were determined for all biocides (Wiegand, Hilpert, and Hancock 2008). The assay was modified by adjusting the concentration ranges and dilution steps and by setting the initial cell concentration to 10^9^ cfu/mL (OD_600_ of 1) because time kill assays were conducted at this density and to account for the inoculation effect (Udekwu et al. 2009). Pre-cultures were adjusted to 10^9^ cfu/mL in 200 µl LB Lennox containing biocide. Cultures were incubated in 96-well plates made from polypropylene (Greiner) for 24 hours at 37°C with shaking and growth was assessed through measurement of the OD in an Epoch plate reader (Biotek). The lowest concentration without observable growth as determined by optical density was designated as the MIC. To determine the MBC, 10 µL samples were taken from at least 3 concentrations above the MIC, serially diluted (dilution of 10^−1^ to 10^−8^) and 10 µL were spotted on LB agar to enumerate colony forming units. The lowest concentration with less than 10 colonies, corresponding to a >10^5^-fold reduction of the initial cell number, was designated the MBC. The concentration ranges used for the individual substances were: H_2_O_2_ [mM]: 0.39, 0.78, 1.56, 3.125, 6.25, 12.5, 25, 50, 100, 200; Glutaraldehyde [%]: 0.02, 0.04, 0.08, 0.16, 0.23, 0.31, 0.47, 0.63, 0.94, 1.25; chlorhexidine [µg/mL]: 2, 3, 4, 5, 6.25, 9.375, 12.5, 18.75, 25, 35; benzalkonium chloride [µg/mL]: 5.3, 7.1, 14.2, 17.7, 21.2, 24.8, 28.3, 31.8, 35.4, 53.1; DDAC [µg/mL]: 14, 16, 24, 32, 40, 48, 56, 64, 72, 80; isopropanol [%]: 0.07, 0.14, 0.275, 0.55, 1.1, 2.2, 4.4, 8.75, 17.5, 35.

### Time kill assays

Time kill assays were carried out in phosphate buffered saline (PBS, final concentrations: 2.68 mM KCl, 1.76 mM KH_2_PO_4_, 10 mM Na_2_PO_4_, 137 mM NaCl) to minimize reactions of the biocides with organic matter in the growth medium. Pre-cultures were pelleted at 4000 g for 4 minutes and the cell density was adjusted to 10^9^ cfu/mL in PBS. Cell density prior to biocide addition was determined by sampling 10 µL, serial dilution (10^−6^-fold) and plating of 100 µL on an LB agar plate. Biocides were added to the desired concentration in a final volume of 900 µL in 2 mL microcentrifuge tubes and samples were incubated in a ThermoMixer at 37°C with agitation at 1200 rpm. Samples of 10 µL were taken at given timepoints, serially diluted in PBS at 10-fold dilution steps and 10 µL of each dilution were spotted on LB agar to enumerate colony forming units. Experiments were performed with 6 replicates. To achieve high temporal resolution, multi-channel pipettes and 96-well plates were employed for sampling and serial dilution as described earlier (Nordholt et al. 2021). The biocide concentrations for the time kill assays were chosen to be in the range of the MBC or higher and such that kill kinetics could still be observed. The concentrations obtained in the MBC assay served as an orientation and the effective, bioactive concentration in the time kill assay was likely higher than in the MBC assay due to quenching by organic matter in LB medium which was used for the MIC and MBC assays. This was especially evident in the case of glutaraldehyde (Table 1).

**Table 1:** Overview of the used substances and their antimicrobial properties.

| Substance<br>[units] | MIC<sup>a</sup> | MBC | Concentration<br>in TKA | Best fit | AIC weight<sup>b</sup><br>(evidence<br>ratio<sup>c</sup>) |
| --- | --- | --- | --- | --- | --- |
| H<sub>2</sub>O<sub>2</sub> [mM] | 12.5 – 25 | 12.5 – 25 | 75 | Single Weibull | 0.81<br>(4.3) |
| GTA [%] | 0.08 – 0.16 | 0.08 – 0.16 | 0.015<sup>d</sup> | Single Weibull | 0.9843<br>(14.945) |
| CHX [µg/ml] | 6.25 – 9.375 | 18.75 – 25 | 100 | Single Weibull | 0.72<br>(2.51) |
| BAC [µg/ml] | 31.8 – 35.4 | 35.4 – 53.1 | 53.1 | Double<br>Weibull | 0.9998<br>(4783.97) |
| DDAC [µg/ml] | 16 – 24 | 16 – 24 | 28 | Double<br>Weibull | 0.999<br>(65659.64) |
| Isopropanol<br>[%] | 2.2 – 4.4 | 4.4 – 8.75 | 12 | Double<br>Weibull | 0.996<br>(257.3) |

<sup>a</sup> The range of assayed concentrations can be found in the methods section. <sup>b</sup> The AIC weight gives the probability of a model to be the best out of all tested models. <sup>c</sup> The evidence ratio of the best model over the second best model. The evidence ratio gives the relative likelihood, i.e. how much more likely the best model is over the second best model. <sup>d</sup> The low concentration in the time-kill assay compared to the MBC for GTA is explained by the absence of organic matter in the time-kill assay buffer.

To test whether resistant mutants were responsible for wide tolerance distributions observed for DDAC and CHX, two colonies originating from cells which survived for 12.5 minutes or longer were picked and subjected to MIC determinations as described above. A colony from a freezer stock served as a control.

### Spiking experiment

To test the possibility that the observed kinetics are due to exhaustion of biocide from the media, a spiking experiment was conducted for CHX (Figure 3). Cells from the original culture were added to an ongoing time kill assay to test whether the multimodal killing kinetics were caused by exhaustion of CHX from the medium. For this, a time kill assay was conducted as described in the main methods, with an initial volume of 900 µl. At 22 minutes, 80 µL of the original culture were added to the remaining 830 µL of the time kill assay (70 µL of were sampled previously). Since the cell concentration in the original culture was known, we calculated the number of cells at time point t = 22 minutes as, on the basis of approx. 10^9^ cfu/ml in the original culture: 0.8×10^7^ cfu / 830 µL × 1000 = 9.64 * 10^7^ cfu/mL. In the following, the time points 22.5, 23, 24, 32 minutes were measured as described above.

### Calculations and Statistics

Equations (1) and (3) were fit to the log-transformed data of colony counts of all biological replicates, using the lmfit package version 1.0.0 for Python 3.8 (Newville et al. 2014). To avoid fitting local optima, an initial fit was done using the ‘differential evolution’ method with randomized initial parameters to approximate the globally optimal parameters. These parameters served as starting point for the ‘least squares’ minimization algorithm. An information theoretical approach was used to rank the models and discriminate between uni- and bimodal distributions, i.e. the absence or presence of a persister subpopulation, respectively. The models were ranked and the most appropriate model was selected based on the Akaike information criterion corrected for small sample size (AICc) (Burnham and Anderson 2002). Akaike weights give the probability of the selected model being the correct one among the tested models. The weights were calculated by dividing the AICc by the sum of the AICc of all models.

The following data points were not considered during analyses: zero counts due to unknown uncertainty in those measurements. Data points which increased more than 3-fold in comparison to the previous timepoint (6 of 316 non-zero data points [1.9%]) and data with only a single replicate measurement (2 of 316 non-zero data points [0.63%]) where not considered due to high chance of resulting from experimental errors.

Equation (3) can be rewritten to obtain a commonly used form of the Weibull distribution. The CDF is then given by:

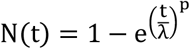

with *p* as the shape parameter as in Eq. (3) and 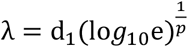, where *d*_*1*_ is the first decimal reduction time as in (3). This definition is equivalent to the `weibull_min()` function in the Python `stats` module.

## Results

### Derivation of Weibull model for chemical inactivation

To model the time-kill kinetics of *E. coli* exposed to lethal concentrations of chlorhexidine (CHX), benzalkonium chloride (BAC), 2-propanol (Iso), didecyldimethylammonium chloride (DDAC), hydrogen peroxide (H_2_O_2_) and glutaraldehyde (GTA), we considered a model based on the survival function of the sum of two mixed Weibull distributions (Coroller et al. 2006). The two distributions describe the tolerance times of two populations, i.e. the major, susceptible population and a minor, more tolerant persister subpopulation. The model describes the number of survivors *N* consisting of the two subpopulations as function of time *t* as:

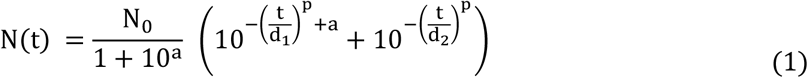

where *N*_*0*_ is the inoculum size in cfu/mL, *a* is the logit transformation of the initial fraction *f* of the susceptible population 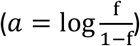 is a shape parameter and *d*_*1*_ and *d*_*2*_ are parameters associated to the tolerance of the populations. Specifically, they are the treatment times for the first decimal reduction of population 1 (susceptible) and 2 (persister), respectively. We use the simplified model by Coroller *et al*. which sets the shape parameter *p* to be the same for both Weibull distributions (Coroller et al. 2006). The shape parameter *p* is related to the heterogeneity in the population via the coefficient of variation (CV = Standard deviation / mean) as follows: if p = 1, CV = 1; if p > 1, CV < 1; and if p < 1, CV > 1. By setting *p* = 1, the model simplifies to the well-known model commonly used in antibiotic persistence research (Balaban et al. 2004), which can also be interpreted as the sum of two exponential distributions:

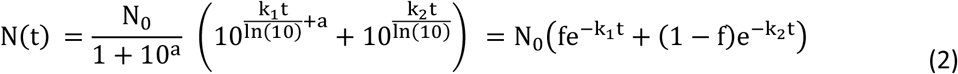

where *k*_*1*_ and *k*_*2*_ are the rate parameters at which each subpopulation dies, with 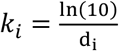. When only one population is present, i.e. fraction *f* = 1, the bimodal equations (1) and (2) simplify to unimodal equations:

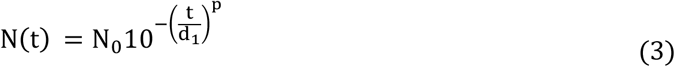

and with

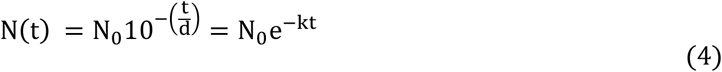

Equation (1), and its special cases Eqs. (2) – (4), can be used to fit a wide range of inactivation kinetics, including those that deviate from the idealistic log-linear decrease, as exemplified in Figure 1. The tolerance time distributions obtained from time kill kinetics are straightforwardly used to sample survival times for statistical modelling approaches (Zafiro Aspridou and Koutsoumanis 2020).

**Figure 1:**
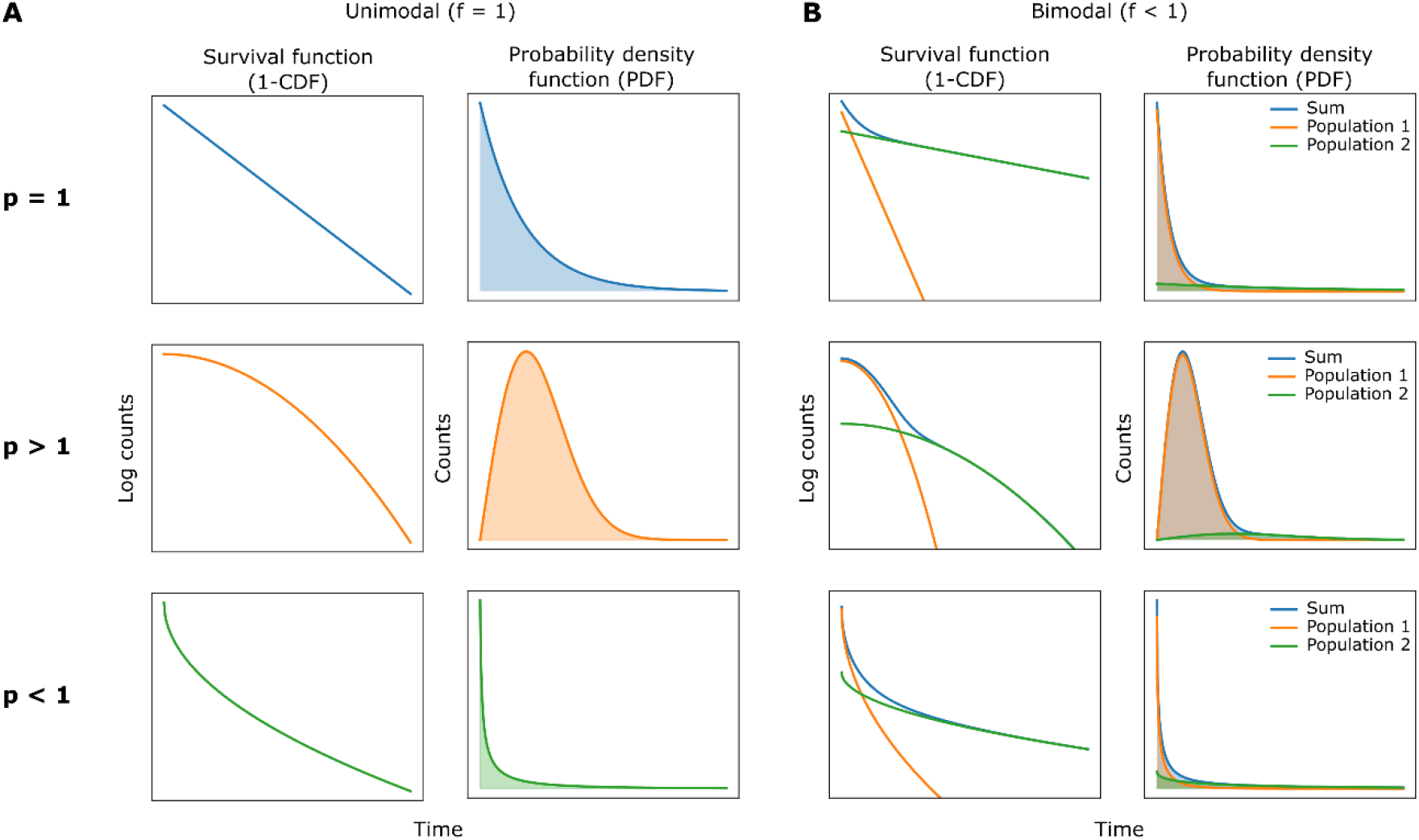
Schematic depiction of different inactivation kinetics as generated by Weibull distributions. Disinfection kinetics can be interpreted as the survival function (left panel) of a Weibull distribution, which can be represented as the probability density function (right panel), highlighting the distribution of individual tolerance times within the bacterial population. **(A)** The shape and the spread of the tolerance time distributions depends on the shape parameter *p*. For the unimodal case *p* = 1 corresponds to a coefficient of variation (CV) = 1; *p* > 1 corresponds to a CV < 1; and *p* < 1 corresponds to a CV > 1. **(B)** Examples of bimodal distributions indicating the presence of two populations, i.e. the fraction *f* of susceptible cells is smaller than 1. The bimodal distribution (blue line) is the sum of two unimodal distributions (orange and green lines) with fractions *f* and 1-*f*, according to equation (1). The shape parameter *p* is the same for the distributions of population 1 and 2, as in (Coroller et al. 2006).

### Disinfection kinetics

The substances that were assayed are used as disinfectants, preservatives and/or antiseptics and belong to the classes of quaternary ammonium compounds (benzalkonium chloride [BAC], DDAC), alcohols (isopropanol), aldehydes (glutaraldehyde [GTA]), oxidative substances (hydrogen peroxide [H_2_O_2_]) and cationic biguanides (chlorhexidine [CHX]). As a first step to characterize their antimicrobial activity, the minimum inhibitory concentration (MIC) and minimum biocidal concentration (MBC) for each substance were determined (Table 1). In most cases, the MBC was in the same concentration range as the MIC. The MIC and MBC determination were carried out with a cell density of 10^9^ cfu/mL, as they were performed to determine the concentrations for the time kill assays to be conducted with an inoculum of 10^9^ cfu/mL and since biocide activity is subject to the inoculum effect, i.e. the antimicrobial activity is affected by the cell density (Udekwu et al. 2009; Loffredo et al. 2021).

The goal of this study was to identify substances against which a population of *E. coli* cells contains persister cells. Therefore, time-kill assays were carried out with populations in stationary phase, which is known to induce persistence (Luidalepp et al. 2011; Balaban et al. 2004; Harms et al. 2017). Bimodal time kill kinetics upon challenge with lethal antimicrobial stress are indicative of a persister subpopulation (Balaban et al. 2019). To distinguish between unimodal and bimodal kill kinetics, equations (1) and (3) were fitted to the data obtained in time kill assays and the models were scored based on the Akaike information criterion (AIC), which takes into account the goodness of fit as well as the parsimony of the model (Burnham and Anderson 2002). The Akaike weights in Table 1 give the probability of the selected model to be the best model under the given data, with the evidence ratio describing how much more likely the selected model was over the alternative model.

The kill kinetics of *E. coli* exposed to lethal biocide concentrations were qualitatively and quantitatively dependent on the identity of the substance used (Figure 2). There was no evidence for the presence of a persister subpopulation against glutaraldehyde and H_2_O_2_ (Figure 2 A, B). Instead, the model suggested that the initial shoulder followed by an accelerating rate of killing in the time-kill curves was due to a unimodal, symmetric distribution of tolerance times with parameter *p* > 1, signifying relatively low heterogeneity within the population (Table 1). In contrast, the disinfection kinetics obtained in the presence of chlorhexidine indicated high heterogeneity of the tolerance times of individual cells (Figure 2 C). However, there was no need to invoke a persister subpopulation to explain the long-tailed time-kill curve (Table 1). Instead, the model suggested that these kinetics were caused by a wide, heavily right-tailed unimodal distribution of tolerance times with shape parameter *p* < 1 (Table 1). While the unimodal model provided the best fit, the evidence ratio over the bimodal model was lower as compared to H_2_O_2_ and GTA (Table 1). Thus, the presence or absence of a persister subpopulation to CHX needs to be investigated in more depth in future studies.

**Figure 2:**
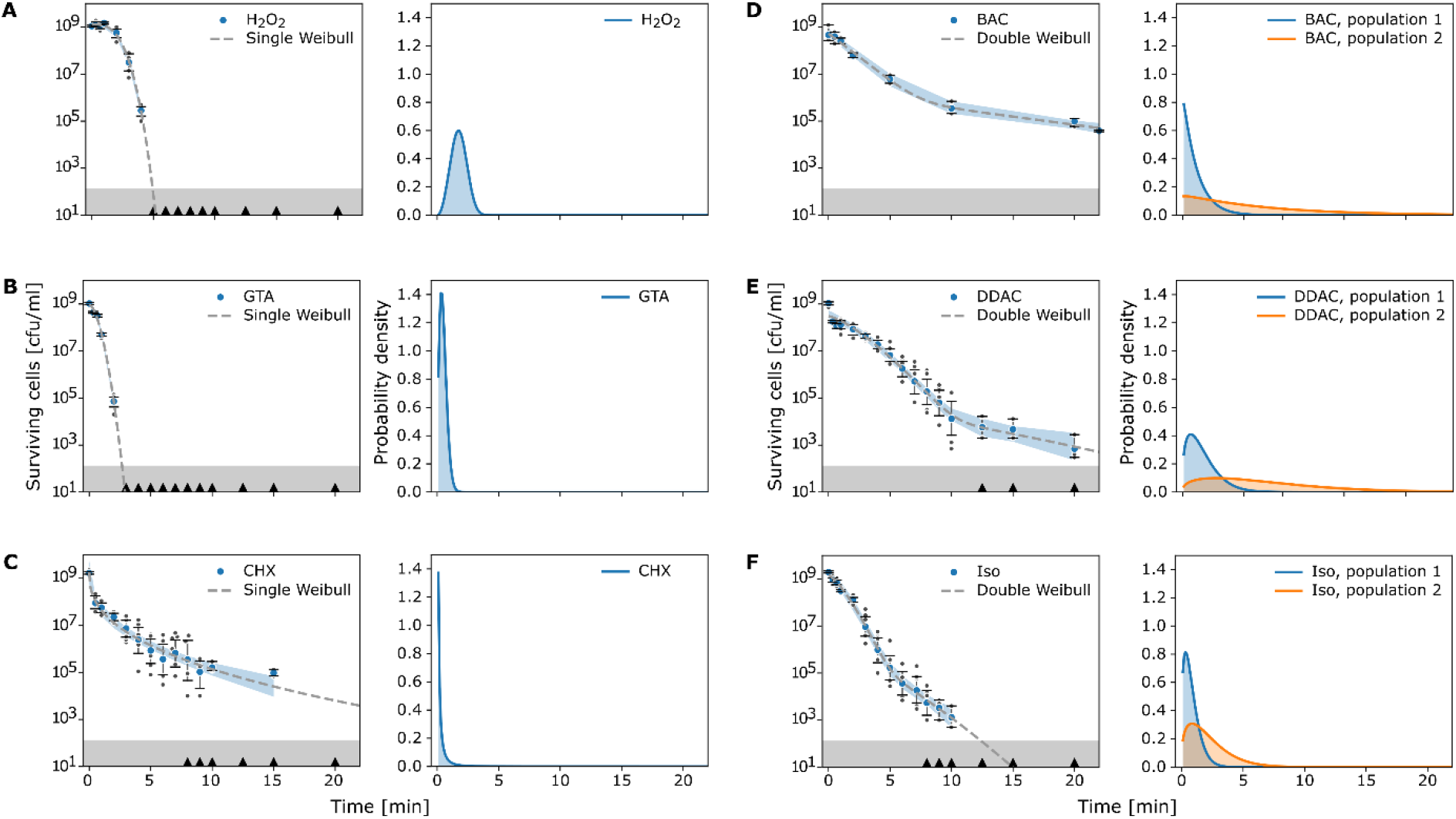
Phenotypic heterogeneity affects disinfection kinetics. Time kill curves of *E. coli* exposed to different disinfectants and Weibull distributions shown as survival functions (left panels) and probability density functions (right panels). Blue circles indicate the geometric mean time kill experiments; error bars indicate the 95% C.I. obtained by bootstrapping. Dark grey circles are datapoints of individual experiments. Black triangles on the x-axes indicate when zero-counts were present. The blue shaded areas indicate the 95 % C.I. of the model fit (dashed line) to the experimental data, excluding values with zero counts. The grey shaded area at the bottom indicates the detection limit. The right panels show the probability density functions of the Weibull distribution(s) fitted to the experimental data (left panel). The blue distribution depicts the major population, the orange distribution shows the persister subpopulation, if present. Number of biological replicates n = 6, except for H_2_O_2_ where n = 5 and BAC where n = 3 biological replicates. **(A)** H_2_O_2_, hydrogen peroxide; **(B)** GTA, glutaraldehyde; **(C)** CHX, chlorhexidine; **(D)** BAC, benzalkonium chloride; **€** DDAC, didecyldimethylammonium chloride; **(F)** Iso, Isopropanol.

**Figure 3:**
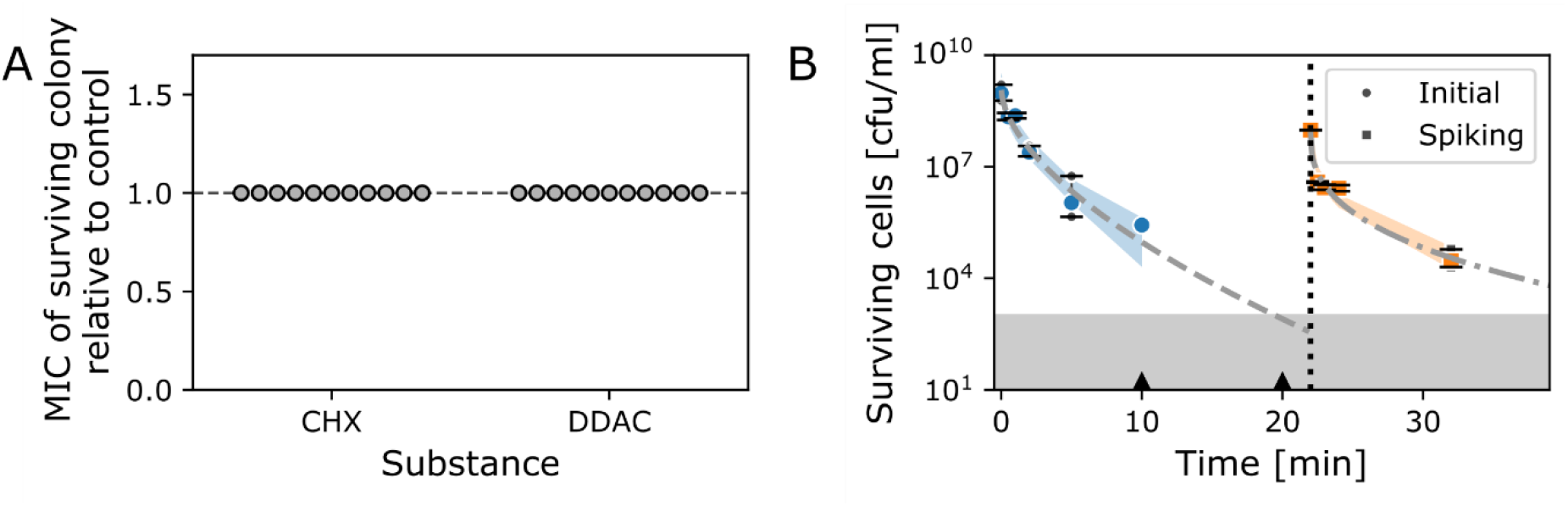
Prolonged time-kill kinetics of CHX and DDAC are not caused by resistance mutations or by exhaustion of biocide from the medium. A) The MIC values of colonies originating from cells which survived a time kill assay with CHX or DDAC for 15 minutes relative to the MIC of a control colony which was not exposed to the biocide. For each biological replicate (1-5), two colonies were picked. One colony was picked for replicate 6 and for the control. The absolute MIC values for each substance are given in Table 1. B) Biological activity of CHX is maintained beyond a 20 minute time kill assay, showing that the time kill kinetics are not caused by exhaustion of CHX. The vertical dotted line indicates the time when cells from the original culture were spiked into the killing assay. Number of biological replicates n = 3. Symbols indicate the geometric mean, error bars indicate the 95% C.I. obtained by bootstrapping. Small grey symbols are datapoints of individual experiments. Black triangles on the x-axis indicate when zero-counts were present. The dashed line and the dash-dotted lines are fits of single Weibull distributions to the data. The shaded areas around the fits indicate the 95 % C.I. of the model fit, excluding values with zero counts. The grey shaded area at the bottom indicates the detection limit.

The other cationic substances, the quaternary ammonium compounds (QACs) DDAC and BAC, also exhibited highly heterogeneous tolerance time distributions (Figure 2 D, E). In these cases, however, there was strong support for the presence of a tolerant persister subpopulation (Table 1). Two alternative hypotheses that could explain long tailed disinfection kinetics were experimentally tested, as was previously done for BAC (Nordholt et al. 2021). First, the possibility of resistance mutations against DDAC and CHX causing the wide tolerance time distributions was excluded by determining that the MIC of cells which survived for 15 minutes remained unchanged (Figure 3 A). Second, we excluded that the loss of activity of the antimicrobial substance over the course of the assay was responsible for the long-tailed kinetics of CHX. To this end, we spiked fresh cells into the late phase of a time kill assay, which again resulted in similar disinfection kinetics (Figure 3 B). These observations corroborate that the disinfection kinetics were caused by the heterogeneous tolerance time distributions within the populations, rather than resistance mutations or changes in the activity of the disinfectant. Likewise, the data obtained from a time kill assay with 12 % isopropanol indicated the presence of a tolerant subpopulation (Figure 2 F). However, the decimal reduction parameter *d*_*2*_ of the second, more tolerant subpopulation was considerably lower than for the quaternary ammonium compounds (Table 2), resulting in an overall faster killing of the population.

**Table 2:** Parameter values of the best fits of the Weibull distribution to the data. Errors indicate the standard error of the estimated parameters.

| Substance | $N_0$ [cfu/ml] | $p$ | $d_1$ [min] | $d_2$ [min] | $a$ |
| --- | --- | --- | --- | --- | --- |
| H<sub>2</sub>O<sub>2</sub> | $1.31 \pm 0.16 \times 10^9$ | $3.03 \pm 0.24$ | $2.60 \pm 0.09$ | n.a. | n.a. |
| GTA | $9.21 \pm 1.27 \times 10^8$ | $1.76 \pm 0.1$ | $0.90 \pm 0.05$ | n.a. | n.a. |
| CHX | $1.8 \pm 1.04 \times 10^9$ | $0.41 \pm 0.06$ | $0.3 \pm 0.17$ | n.a. | n.a. |
| BAC | $5.16 \pm 1.88 \times 10^8$ | $1.03 \pm 0.20$ | $2.52 \pm 0.44$ | $14.99 \pm 3.72$ | $2.52 \pm 0.29$ |
| DDAC | $3.02 \pm 0.85 \times 10^8$ | $1.35 \pm 0.16$ | $3.32 \pm 0.4$ | $13.69 \pm 4.65$ | $3.87 \pm 0.58$ |
| Isopropanol | $1.47 \pm 0.51 \times 10^9$ | $1.33 \pm 0.21$ | $1.68 \pm 0.27$ | $4.45 \pm 0.98$ | $3.12 \pm 0.48$ |

## Discussion

The model which was used here to describe the time kill kinetics has interesting properties, which make it an attractive tool in the interpretation of time kill kinetics. From a conceptual point of view, the interpretation of time kill curves as probability distributions highlights the heterogeneity of individual cells. This means that time kill kinetics can be described by the properties of the distributions, i.e. the moments, and the quantiles of the distribution. The obtained distributions can readily be used for sampling individual cells in stochastic modelling approaches (Zafiro Aspridou and Koutsoumanis 2020). Another strength of the model is its flexibility with regard to the shapes of time kill curves that can be modelled.

In the interpretation of the model parameters, it is instructive to relate our findings for chemical inactivation to the findings of Coroller *et al*. in their physical inactivation studies. In their study on the effect of growth stage on the tolerance to physical inactivation by heat, Coroller *et al*. studied the evolution of the model parameters over the course of bacterial growth curves. The external stress was kept constant, whereas the physiology of the bacterial cultures varied. They found that the decimal reduction parameter of the susceptible population, *d*_*1*_, as well as the fraction of tolerant cells increased with entry into stationary phase, whereas the decimal reduction parameter of the tolerant subpopulation, *d*_*2*_, and the shape parameter *p* were largely independent of the physiological state of the cells. In contrast, our experimental design kept the physiological state of the cells constant and varied the external stress. We found that all model parameters were dependent on the substance class of the disinfectant (Table 2). It can be speculated that the parameter *p* is related to the interaction between tolerance mechanisms and the mode(s) of action of the tested substance, suggesting that *p* for a given bacterial strain depends on the substance. In our recent study on BAC, we found that *d*_*1*_ and the fraction of the tolerant subpopulation were concentration dependent, whereas *d*_*2*_ appeared to be largely independent of the BAC concentration (Nordholt et al. 2021). Similar observations were made previously (García and Cabo 2018; Imlay and Linn 1986). Interestingly, the parameters obtained here for the quaternary ammonium compounds BAC and DDAC were quite similar, likely due to a similar mode of action (Table 2).

The wide distributions observed for the cationic agents CHX, BAC and DDAC imply high heterogeneity in the ability of individual *E. coli* cell to tolerate these substances. Recently, Şimşek and Kim showed that the distribution of a tolerance trait, namely the lag time, can determine antibiotic killing kinetics (Şimşek and Kim 2019). It is possible that the kinetics which we observed here are shaped by the distributions of one or more traits which affect tolerance. There are several traits which are known to affect susceptibility to cationic antimicrobials. For example, cationic substances are thought to mainly act on the bacterial membrane, which bears a net-negative charge (Gilbert and Moore 2005). Thus, phenotypic differences in membrane composition could endow individual bacteria with a higher tolerance, for example through changes to the surface charge or membrane permeability. In support of this idea are several reports that found genetic alterations pertaining membrane components allow the adaptation to cationic antimicrobials (Nordholt et al. 2021; Nagai et al. 2003; Wand et al. 2017; Kim et al. 2018; Herrera, Hankins, and Trent 2010; Tag ElDein et al. 2021). Another general defense mechanism against QACs and CHX is multi-drug efflux (Buffet-Bataillon et al. 2012; Gregorchuk et al. 2021; Nordholt et al. 2021). Efflux pump content has been shown to be heterogeneous in isogenic populations (Bergmiller et al. 2017; Pu et al. 2016), and so it is possible that heterogeneous efflux activity contributes to the wide tolerance time distributions observed for the cationic antimicrobials. In addtion, synergistic interactions between multiple tolerance traits in persister subpopulations are conceivable. Measuring the heterogeneity in these traits and correlating them with killing kinetics could be informative as to whether they underlie the wide tolerance distributions. However, it should be noted that persister subpopulations themselves appear to be highly heterogeneous, with their defining and unifying trait being increased tolerance (Goormaghtigh and Van Melderen 2019; Balaban et al. 2019; Şimşek and Kim 2019), which indicates that the tolerance mechanisms are at least partially substance specific and likely multifactorial.

Despite their similar mode of action (Gilbert and Moore 2005), the specificity of efflux pumps and the adaptive mechanisms to CHX and QACs can be quite different (Slipski, Zhanel, and Bay 2018; Merchel Piovesan Pereira, Wang, and Tagkopoulos 2021). This might explain why a tolerant subpopulation was detected against QACs, but not against CHX. Certain regulatory motifs or stochastic events can promote heterogeneity and result in multimodal distributions of expression levels in individual cells (Choi et al. 2008; Thattai and van Oudenaarden 2001; Dubnau and Losick 2006), meaning that the identity and the regulation of a specific tolerance trait could underlie the presence of a persister subpopulation. For example, in our recent study on BAC tolerance, we found that changes in the outer membrane charge reduce the susceptibility to the cationic BAC, but not to the cationic antibiotic colistin, suggesting that the underlying tolerance mechanisms are indeed substance specific (Nordholt et al. 2021; Herrera, Hankins, and Trent 2010).

Our knowledge on mechanisms that increase tolerance to biocides is sparse compared to the mechanisms that confer resistance, as measured by an increase in the MIC. From antibiotic research it is known that the mechanisms that increase tolerance and mechanisms that increase resistance do not necessarily overlap (Brauner et al. 2016; Van den Bergh et al. 2016; Levin-Reisman et al. 2017) and our recent findings indicate that this could also be the case for biocides (Nordholt et al. 2021).

Wide disinfection kinetics are potentially problematic for two reasons: first, treatment times could be underestimated, resulting in survival of bacteria in the tail of the distribution. Second, it can be argued that high phenotypic heterogeneity in the tolerance of individual cells implies a high potential for genetic adaptation of the underlying trait when cells are selected for their tolerance. Although increased tolerance is their defining trait, persister subpopulations often display properties that set them apart from the majority of the population, such as dormancy or low metabolic activity, contributing to their perception as distinct subpopulations (Pu et al. 2018; Radzikowski et al. 2016). What could be the consequences when a persister subpopulation is present during disinfection? Recent studies suggest that persister subpopulations facilitate the evolution of resistance by other means than merely ensuring the survival of the population, for example through their propensity for acquiring mutations (El Meouche and Dunlop 2018; Windels et al. 2019; Salini et al. 2022). In line with this, we recently reported that persistence can facilitate the rapid evolution of tolerance against BAC (Nordholt et al. 2021). Whether this can be generalized to other substances that exhibit multimodal kill kinetics, such as DDAC and Isopropanol, and whether adaption in the presence of persisters occurs faster than when cells exhibit a wide unimodal distribution, as observed for CHX, remains to be shown, for example through experimental evolution. Here we provide a conceptual and analytical framework as a starting point for such studies.

## References

Ackermann, Martin. 2015. “A Functional Perspective on Phenotypic Heterogeneity in Microorganisms.” Nature Publishing Group 13 (8): 497–508. https://doi.org/10.1038/nrmicro3491.

Aspridou, Zafeiro, Athanasios Balomenos, Panagiotis Tsakanikas, Elias Manolakos, and Konstantinos Koutsoumanis. 2019. “Heterogeneity of Single Cell Inactivation: Assessment of the Individual Cell Time to Death and Implications in Population Behavior.” Food Microbiology 80 (June): 85–92. https://doi.org/10.1016/j.fm.2018.12.011.

Aspridou, Zafiro, and Konstantinos Koutsoumanis. 2020. “Variability in Microbial Inactivation: From Deterministic Bigelow Model to Probability Distribution of Single Cell Inactivation Times.” Food Research International 137 (November): 109579. https://doi.org/10.1016/j.foodres.2020.109579.

Balaban, Nathalie Q., Sophie Helaine, Kim Lewis, Martin Ackermann, Bree Aldridge, Dan I. Andersson, Mark P. Brynildsen, et al. 2019. “Definitions and Guidelines for Research on Antibiotic Persistence.” Nature Reviews Microbiology, April, 1. https://doi.org/10.1038/s41579-019-0196-3.

Balaban, Nathalie Q., Jack Merrin, Remy Chait, Lukasz Kowalik, and Stanislas Leibler. 2004. “Bacterial Persistence as a Phenotypic Switch.” Science 305 (5690). http://science.sciencemag.org/content/305/5690/1622.long.

Bergmiller, Tobias, Anna M. C. Andersson, Kathrin Tomasek, Enrique Balleza, Daniel J. Kiviet, Robert Hauschild, GaŠper TkaČik, and Călin C. Guet. 2017. “Biased Partitioning of the Multidrug Efflux Pump AcrAB-TolC Underlies Long-Lived Phenotypic Heterogeneity.” Science 356 (6335): 311–15. https://doi.org/10.1126/science.aaf4762.

Blattner, F. R., G. Plunkett, C. A. Bloch, N. T. Perna, V. Burland, M. Riley, J. Collado-Vides, et al. 1997. “The Complete Genome Sequence of Escherichia Coli K-12.” Science (New York, N.Y.) 277 (5331): 1453–62. https://doi.org/10.1126/science.277.5331.1453.

Bore, Erlend, Michel Hébraud, Ingrid Chafsey, Christophe Chambon, Camilla Skjæret, Birgitte Moen, Trond Møretrø, Øyvind Langsrud, Knut Rudi, and Solveig Langsrud. 2007. “Adapted Tolerance to Benzalkonium Chloride in Escherichia Coli K-12 Studied by Transcriptome and Proteome Analyses.” Microbiology 153 (4): 935–46. https://doi.org/10.1099/mic.0.29288-0.

Boxtel, Coco van, Johan H. van Heerden, Niclas Nordholt, Phillipp Schmidt, and Frank J Bruggeman. 2017. “Taking Chances and Making Mistakes: Non-Genetic Phenotypic Heterogeneity and Its Consequences for Surviving in Dynamic Environments.” Journal of The Royal Society Interface 14 (132). https://doi.org/10.1098/rsif.2017.0141.

Brauner, Asher, Ofer Fridman, Orit Gefen, and Nathalie Q. Balaban. 2016. “Distinguishing between Resistance, Tolerance and Persistence to Antibiotic Treatment.” Nature Reviews Microbiology 14 (5): 320–30. https://doi.org/10.1038/nrmicro.2016.34.

Buffet-Bataillon, Sylvie, André Le jeune, Sandrine Le Gall-David, Martine Bonnaure-Mallet, and Anne Jolivet-Gougeon. 2012. “Molecular Mechanisms of Higher MICs of Antibiotics and Quaternary Ammonium Compounds for Escherichia Coli Isolated from Bacteraemia.” Journal of Antimicrobial Chemotherapy 67 (12): 2837–42. https://doi.org/10.1093/jac/dks321.

Burnham, K.P., and D.R. Anderson. 2002. Model Selection and Multimodel Inference: A Practical Information-Theoretic Approach (2nd Ed). Vol. 172. http://linkinghub.elsevier.com/retrieve/pii/S0304380003004526.

Cameron, David R, Yue Shan, Eliza A Zalis, Vincent Isabella, and Kim Lewis. 2018. “A Genetic Determinant of Persister Cell Formation in Bacterial Pathogens.” Journal of Bacteriology, June, JB.00303-18. https://doi.org/10.1128/JB.00303-18.

Choi, Paul J., Long Cai, Kirsten Frieda, and X. Sunney Xie. 2008. “A Stochastic Single-Molecule Event Triggers Phenotype Switching of a Bacterial Cell.” Science 322 (5900): 442–46. https://doi.org/10.1126/science.1161427.

Coroller, L., I. Leguerinel, E. Mettler, N. Savy, and P. Mafart. 2006. “General Model, Based on Two Mixed Weibull Distributions of Bacterial Resistance, for Describing Various Shapes of Inactivation Curves.” Applied and Environmental Microbiology 72 (10): 6493–6502. https://doi.org/10.1128/AEM.00876-06.

Dörr, Tobias, Kim Lewis, and Marin Vulić. 2009. “SOS Response Induces Persistence to Fluoroquinolones in Escherichia Coli.” Edited by Susan M. Rosenberg. PLoS Genetics 5 (12): e1000760. https://doi.org/10.1371/journal.pgen.1000760.

Dubnau, David, and Richard Losick. 2006. “Bistability in Bacteria.” Molecular Microbiology 61 (3): 564–72. https://doi.org/10.1111/j.1365-2958.2006.05249.x.

El Meouche, Imane El, and Mary J. Dunlop. 2018. “Heterogeneity in Efflux Pump Expression Predisposes Antibiotic-Resistant Cells to Mutation.” Science 362 (6415): 686–90. https://doi.org/10.1126/science.aar7981.

European Commission Environment Directorate-General. 2009. “Assessment of Different Options to Address Risks from the Use Phase of Biocides - Final Report.” 4. European Commission Environment Directorate-General. https://ec.europa.eu/environment/archives/ppps/pdf/final_report0309.pdf.

García, Míriam R, and Marta L Cabo. 2018. “Optimization of E. Coli Inactivation by Benzalkonium Chloride Reveals the Importance of Quantifying the Inoculum Effect on Chemical Disinfection.” Frontiers in Microbiology 9 (JUN): 1259. https://doi.org/10.3389/fmicb.2018.01259.

Gilbert, Peter, and L. E. Moore. 2005. “Cationic Antiseptics: Diversity of Action under a Common Epithet.” Journal of Applied Microbiology 99 (4): 703–15. https://doi.org/10.1111/j.1365-2672.2005.02664.x.

Goormaghtigh, Frédéric, and Laurence Van Melderen. 2019. “Single-Cell Imaging and Characterization of Escherichia Coli Persister Cells to Ofloxacin in Exponential Cultures.” Science Advances 5 (6): eaav9462. https://doi.org/10.1126/sciadv.aav9462.

Gregorchuk, Branden S. J., Shelby L. Reimer, Kari A. C. Green, Nicola H. Cartwright, Daniel R. Beniac, Shannon L. Hiebert, Timothy F. Booth, et al. 2021. “Phenotypic and Multi-Omics Characterization of Escherichia Coli K-12 Adapted to Chlorhexidine Identifies the Role of MlaA and Other Cell Envelope Alterations Regulated by Stress Inducible Pathways in CHX Resistance.” Frontiers in Molecular Biosciences 8. https://doi.org/10.3389/fmolb.2021.659058.

Harms, Alexander, Cinzia Fino, Michael A Sørensen, Szabolcs Semsey, and Kenn Gerdes. 2017. “Prophages and Growth Dynamics Confound Experimental Results with Antibiotic-Tolerant Persister Cells.” MBio 8 (6): e01964–17. https://doi.org/10.1128/mBio.01964-17.

Herrera, Carmen M., Jessica V. Hankins, and M. Stephen Trent. 2010. “Activation of PmrA Inhibits LpxT-Dependent Phosphorylation of Lipid A Promoting Resistance to Antimicrobial Peptides.” Molecular Microbiology 76 (6): 1444–60. https://doi.org/10.1111/j.1365-2958.2010.07150.x.

Imlay, J A, and S Linn. 1986. “Bimodal Pattern of Killing of DNA-Repair-Defective or Anoxically Grown Escherichia Coli by Hydrogen Peroxide.” Journal of Bacteriology 166 (2): 519–27. https://doi.org/10.1128/jb.166.2.519-527.1986.

Kampf, Günter. 2018. “Biocidal Agents Used for Disinfection Can Enhance Antibiotic Resistance in Gram-Negative Species.” Antibiotics 7 (4). https://doi.org/10.3390/antibiotics7040110.

Kim, Minjae, Janet K. Hatt, Michael R. Weigand, Raj Krishnan, Spyros G. Pavlostathis, and Konstantinos T. Konstantinidis. 2018. “Genomic and Transcriptomic Insights into How Bacteria Withstand High Concentrations of Benzalkonium Chloride Biocides.” Applied and Environmental Microbiology 84 (12). https://doi.org/10.1128/AEM.00197-18.

Levin-Reisman, Irit, Irine Ronin, Orit Gefen, Ilan Braniss, Noam Shoresh, and Nathalie Q. Balaban. 2017. “Antibiotic Tolerance Facilitates the Evolution of Resistance.” Science 355 (6327): 826– 30. https://doi.org/10.1126/science.aaj2191.

Loffredo, Maria Rosa, Filippo Savini, Sara Bobone, Bruno Casciaro, Henrik Franzyk, Maria Luisa Mangoni, and Lorenzo Stella. 2021. “Inoculum Effect of Antimicrobial Peptides.” Proceedings of the National Academy of Sciences of the United States of America 118 (21): e2014364118. https://doi.org/10.1073/pnas.2014364118.

Luidalepp, Hannes, Arvi Jõers, Niilo Kaldalu, and Tanel Tenson. 2011. “Age of Inoculum Strongly Influences Persister Frequency and Can Mask Effects of Mutations Implicated in Altered Persistence.” Journal of Bacteriology 193 (14): 3598–3605. https://doi.org/10.1128/JB.00085-11.

Merchel Piovesan Pereira Beatriz, Xiaokang Wang, and Ilias Tagkopoulos. 2021. “Biocide-Induced Emergence of Antibiotic Resistance in Escherichia Coli.” Frontiers in Microbiology 12. https://doi.org/10.3389/fmicb.2021.640923.

Murray, Christopher JL, Kevin Shunji Ikuta, Fablina Sharara, Lucien Swetschinski, Gisela Robles Aguilar, Authia Gray, Chieh Han, et al. 2022. “Global Burden of Bacterial Antimicrobial Resistance in 2019: A Systematic Analysis.” The Lancet 0 (0). https://doi.org/10.1016/S0140-6736(21)02724-0.

Nagai, Katsuhiro, Takahiro Murata, Shin Ohta, Hiroshi Zenda, Makoto Ohnishi, and Tetsuya Hayashi. 2003. “Two Different Mechanisms Are Involved in the Extremely High-Level Benzalkonium Chloride Resistance of a Pseudomonas Fluorescens Strain.” Microbiology and Immunology 47 (10): 709–15. https://doi.org/10.1111/j.1348-0421.2003.tb03440.x.

Newville, Matthew, Till Stensitzki, Daniel B. Allen, and Antonino Ingargiola. 2014. LMFIT: Non-Linear Least-Square Minimization and Curve-Fitting for Python. Zenodo. https://doi.org/10.5281/zenodo.11813.

Nordholt, Niclas, Orestis Kanaris, Selina B. I. Schmidt, and Frank Schreiber. 2021. “Persistence against Benzalkonium Chloride Promotes Rapid Evolution of Tolerance during Periodic Disinfection.” Nature Communications 12 (1): 6792. https://doi.org/10.1038/s41467-021-27019-8.

Peleg, Micha, and Martin B. Cole. 1998. “Reinterpretation of Microbial Survival Curves.” Critical Reviews in Food Science and Nutrition 38 (5): 353–80. https://doi.org/10.1080/10408699891274246.

Pu, Yingying, Yingxing Li, Xin Jin, Tian Tian, Qi Ma, Ziyi Zhao, Ssu-yuan Lin, et al. 2018. “ATP-Dependent Dynamic Protein Aggregation Regulates Bacterial Dormancy Depth Critical for Antibiotic Tolerance.” Molecular Cell 73 (1): 143-156.e4. https://doi.org/10.1016/j.molcel.2018.10.022.

Pu, Yingying, Zhilun Zhao, Yingxing Li, Jin Zou, Qi Ma, Yanna Zhao, Yuehua Ke, et al. 2016. “Enhanced Efflux Activity Facilitates Drug Tolerance in Dormant Bacterial Cells.” Molecular Cell 62 (2): 284–94. https://doi.org/10.1016/j.molcel.2016.03.035.

Radzikowski, Jakub Leszek, Silke Vedelaar, David Siegel, Álvaro Dario Ortega, Alexander Schmidt, and Matthias Heinemann. 2016. “Bacterial Persistence Is an Active ΣS Stress Response to Metabolic Flux Limitation.” Molecular Systems Biology 12 (9): 882. https://doi.org/10.15252/msb.20166998.

Ronin, Irine, Naama Katsowich, Ilan Rosenshine, and Nathalie Q Balaban. 2017. “A Long-Term Epigenetic Memory Switch Controls Bacterial Virulence Bimodality.” ELife 6 (February): e19599. https://doi.org/10.7554/eLife.19599.

Salini, S., Sinchana G. Bhat, Saba Naz, Ramanathan Natesh, R. Ajay Kumar, Vinay Kumar Nandicoori, and Krishna Kurthkoti. 2022. “The Error-Prone Polymerase DnaE2 Mediates the Evolution of Antibiotic Resistance in Persister Mycobacterial Cells.” Antimicrobial Agents and Chemotherapy 66 (3): e0177321. https://doi.org/10.1128/AAC.01773-21.

SCENIHR. 2009. “Assessment of the Antibiotic Resistance Effects of Biocides,” no. January: 118. https://doi.org/10.2772/8624.

Schreiber, Frank, and Martin Ackermann. 2020. “Environmental Drivers of Metabolic Heterogeneity in Clonal Microbial Populations.” Current Opinion in Biotechnology 62 (April): 202–11. https://doi.org/10.1016/j.copbio.2019.11.018.

Schreiber, Frank, Sten Littmann, Gaute Lavik, Stéphane Escrig, Anders Meibom, M.M.M. Kuypers, and Martin Ackermann. 2016. “Phenotypic Heterogeneity Driven by Nutrient Limitation Promotes Growth in Fluctuating Environments.” Nature Microbiology in press (May): 1–7. https://doi.org/10.1038/nmicrobiol.2016.55.

Şimşek, Emrah, and Minsu Kim. 2019. “Power-Law Tail in Lag Time Distribution Underlies Bacterial Persistence.” Proceedings of the National Academy of Sciences 116 (36): 17635–40. https://doi.org/10.1073/pnas.1903836116.

Slipski, Carmine J, George G Zhanel, and Denice C Bay. 2018. “Biocide Selective TolC-Independent Efflux Pumps in Enterobacteriaceae.” Journal of Membrane Biology 251 (1): 15–33. https://doi.org/10.1007/s00232-017-9992-8.

Tag ElDein, Moustafa A., Aymen S. Yassin, Ossama El-Tayeb, and Mona T. Kashef. 2021. “Chlorhexidine Leads to the Evolution of Antibiotic-Resistant Pseudomonas Aeruginosa.” European Journal of Clinical Microbiology & Infectious Diseases, June. https://doi.org/10.1007/s10096-021-04292-5.

Thattai, M, and A van Oudenaarden. 2001. “Intrinsic Noise in Gene Regulatory Networks.” Proceedings of the National Academy of Sciences of the United States of America 98 (15): 8614–19. https://doi.org/10.1073/pnas.151588598.

Udekwu, Klas I, Nicholas Parrish, Peter Ankomah, Fernando Baquero, and Bruce R Levin. 2009. “Functional Relationship between Bacterial Cell Density and the Efficacy of Antibiotics.” Journal of Antimicrobial Chemotherapy 63 (4): 745–57. https://doi.org/10.1093/jac/dkn554.

Van den Bergh, Bram, Joran E. Michiels, Tom Wenseleers, Etthel M. Windels, Pieterjan Vanden Boer, Donaat Kestemont, Luc De Meester, et al. 2016. “Frequency of Antibiotic Application Drives Rapid Evolutionary Adaptation of Escherichia Coli Persistence.” Nature Microbiology 1 (5): 16020. https://doi.org/10.1038/nmicrobiol.2016.20.

Van Heerden, Johan H., Hermannus Kempe, Anne Doerr, Timo Maarleveld, Niclas Nordholt, and Frank J. Bruggeman. 2017. “Statistics and Simulation of Growth of Single Bacterial Cells: Illustrations with B. Subtilis and E. Coli.” Scientific Reports 7 (1): 16094. https://doi.org/10.1038/s41598-017-15895-4.

Wand, Matthew E., Lucy J. Bock, Laura C. Bonney, and J. Mark Sutton. 2017. “Mechanisms of Increased Resistance to Chlorhexidine and Cross-Resistance to Colistin Following Exposure of Klebsiella Pneumoniae Clinical Isolates to Chlorhexidine.” Antimicrobial Agents and Chemotherapy 61 (1): e01162–16. https://doi.org/10.1128/AAC.01162-16.

Wiegand, Irith, Kai Hilpert, and Robert E W Hancock. 2008. “Agar and Broth Dilution Methods to Determine the Minimal Inhibitory Concentration (MIC) of Antimicrobial Substances.” Nature Protocols 3 (2): 163–75. https://doi.org/10.1038/nprot.2007.521.

Windels, Etthel Martha, Joran Elie Michiels, Maarten Fauvart, Tom Wenseleers, Bram Van den Bergh, and Jan Michiels. 2019. “Bacterial Persistence Promotes the Evolution of Antibiotic Resistance by Increasing Survival and Mutation Rates.” ISME Journal, no. January. https://doi.org/10.1038/s41396-019-0344-9.

Withell, E. R. 1942. “The Significance of the Variation in Shape of Time-Survivor Curves.” The Journal of Hygiene 42 (2): 124–83.

